# Heterogeneity of EEG resting-state brain networks in absolute pitch

**DOI:** 10.1101/2020.05.03.063206

**Authors:** Marielle Greber, Carina Klein, Simon Leipold, Silvano Sele, Lutz Jäncke

## Abstract

The neural basis of absolute pitch (AP), the ability to effortlessly identify a musical tone without an external reference, is poorly understood. One of the key questions is whether perceptual or cognitive processes underlie the phenomenon as both sensory and higher-order brain regions have been associated with AP. One approach to elucidate the neural underpinnings of a specific expertise is the examination of resting-state networks.

Thus, in this paper, we report a comprehensive functional network analysis of intracranial resting-state EEG data in a large sample of AP musicians (n = 54) and non-AP musicians (n = 51). We adopted two analysis approaches: First, we applied an ROI-based analysis to examine the connectivity between the auditory cortex and the dorsolateral prefrontal cortex (DLPFC) using several established functional connectivity measures. This analysis is a replication of a previous study which reported increased connectivity between these two regions in AP musicians. Second, we performed a whole-brain network-based analysis on the same functional connectivity measures to gain a more complete picture of the brain regions involved in a possibly large-scale network supporting AP ability.

In our sample, the ROI-based analysis did not provide evidence for an AP-specific connectivity increase between the auditory cortex and the DLPFC. In contrast, the whole-brain analysis revealed three networks with increased connectivity in AP musicians comprising nodes in frontal, temporal, subcortical, and occipital areas. Commonalities of the networks were found in both sensory and higher-order brain regions of the perisylvian area. Further research will be needed to confirm these exploratory results.

## Introduction

Absolute pitch (AP) is the rare ability to effortlessly identify the pitch of a musical tone without the aid of an external reference tone (Deutsch, 2013). The neural mechanisms underlying AP are poorly understood. One central issue concerns the question of to what extent perceptual and cognitive processes contribute to the phenomenon. On the one hand, evidence from both structural and functional neuroimaging points towards an involvement of auditory regions (Schlaug *et al*., 1995; Keenan *et al*., 2001; McKetton *et al*., 2019), supporting the view of altered perceptual processing in AP. On the other hand, the two-component model, a prominent cognitive theory of AP, postulates that the association of long-term pitch representations with their labels (pitch labeling) constitutes the neurophysiological fundament of AP (Levitin, 1994). This pitch-labeling process has been associated with neural activation in the dorsolateral prefrontal cortex (DLPFC) (Zatorre *et al*., 1998; Bermudez & Zatorre, 2005).

Aiming to integrate the perceptual and cognitive perspectives on AP, the current study examined EEG resting-state connectivity for contributions of both sensory and higher-order brain regions to AP networks. Electroencephalographic resting-state activity has repeatedly been demonstrated to contain stable individual-specific information (e.g., Paranjape *et al*., 2001; Poulos *et al*., 2002; Näpflin *et al*., 2007; Valizadeh *et al*., 2019). Additionally, it has been shown that music-specific networks can be observed during resting state: Professional musicians exhibit increased EEG resting-state connectivity between brain regions that are involved in music processing and music production (Klein *et al*., 2015). Resting-state connectivity patterns in AP musicians might similarly reflect a network of brain regions underlying this specific expertise.

Analyzing resting-state EEG, a previous study of our group (Elmer *et al*., 2015) found some evidence that the auditory cortex and the DLPFC in the left hemisphere were functionally more strongly connected in AP musicians than in non-AP musicians. However, the study focused solely on these two regions of interest (ROIs) within each hemisphere. While this ROI-based approach minimizes the multiple comparisons problem, it neglects the possibility that the two ROIs could be part of a more extensive network. According to current scientific knowledge, various cognitive functions rely on interactions between distributed brain regions organized within large-scale networks (Sporns *et al*., 2004; Fuster, 2006; Bressler & Menon, 2010; Petersen & Sporns, 2015). The same might apply to AP. Findings from fMRI resting-state studies are in line with a more widespread resting-state network in AP musicians. A graph-theoretical study revealed increased clustering, degrees, strength, and local efficiency during rest in AP musicians not only in the superior temporal gyrus but also on a whole-brain level (Loui *et al*., 2012). Another fMRI study reported increased resting-state connectivity between the right planum polare and the auditory cortex (Kim & Knösche, 2017a). More recently, Brauchli and colleagues (2019a) identified increased local resting-state functional connectivity in the left anterior middle frontal gyrus (in the vicinity of the DLPFC) and in the left intraparietal sulcus, and increased global resting-state functional connectivity in the right superior parietal lobule. This suggests an AP-specific network in higher-order cognitive areas. However, when applying multivariate pattern analysis (MVPA), which can capture more fine-grained connectivity patterns, the classification accuracy for AP and non-AP musicians was highest in the left Heschl’s gyrus.

Taken together, AP-specific resting-state networks may rely on additional temporo-frontal connections besides the one between the auditory cortex and the DLPFC. Whole-brain analyses provide an opportunity to explore this potential involvement of other regions in an AP-specific network. On the downside, in case of stringent multiple testing correction, whole-brain analyses may miss regional connectivity differences that could have been picked up by ROI-based analyses.

A common limitation of most previous neuroscientific studies comparing AP and non-AP musicians are small sample sizes. This is mostly due to the low prevalence of AP as well as the resource-intensive data acquisition in neuroimaging. Small samples result in low statistical power and unreliable estimates of the true effect (Button *et al*., 2013). Therefore, studies with larger samples are urgently needed to advance our understanding of the neural underpinnings of AP.

Using a large sample of musicians with AP (n = 54) and without AP (n = 51), we here reevaluate the question of whether AP musicians demonstrate specific functional resting-state connectivity patterns. We recorded resting-state EEG and employed well-established source estimation techniques to measure functional connectivity. For AP research, EEG-based measures might be particularly suited to estimate neurophysiological coactivations during rest since, in contrast to resting-state fMRI recordings, the data is acquired in silence without background noise. The current study further benefits from the application of several connectivity measures (lagged phase synchronization, lagged linear connectivity, and instantaneous linear connectivity), which are each associated with different strengths and weaknesses regarding volume conduction, individual-specific stability, and relation to structural connectivity as described in detail below (see section on ‘EEG Source-Level Connectivity’ in ‘Material and Methods’).

To combine the methodological advantages of both ROI-based and whole-brain analyses, we adopted two approaches: (1) We conducted an ROI-based analysis to examine the functional connectivity between the auditory cortex and the DLPFC. This part of the study is a replication of the above-described previous study of our group (Elmer *et al*., 2015), which had a much smaller sample. (2) We conducted a whole-brain connectivity analysis to explore a potential involvement of other regions besides the auditory cortex and the DLPFC with regard to a more widespread AP-specific network. This analysis was guided by the findings discussed above, which suggest distributed network features in AP musicians comprising brain areas other than the auditory cortex and the DLPFC.

## Material and Methods

### Participants

Fifty-four AP musicians and 51 non-AP musicians aged 18 – 44 years participated in the EEG resting-state study. All participants were professional musicians, music students, or highly-trained amateur musicians, who were recruited within a larger research project investigating the neural correlates of AP (Greber *et al*., 2018; Brauchli *et al*., 2019a, 2019b; Burkhard *et al*., 2019, 2020; Leipold, Brauchli, Greber, & Jäncke, 2019; Leipold, Greber, Sele, *et al*., 2019; Leipold, Oderbolz, Greber, & Jäncke, 2019). The participants were assigned to the two groups based on self-report. Before being invited to the study, participants underwent online testing assessing demographic information, musical experience, and pitch-labeling ability. Based on these data, the two groups were matched for sex, age, handedness, age of onset of musical training, and cumulative hours of musical training over the lifespan.

None of the participants reported any audiological, neurological, or severe psychiatric disorders. Pure-tone audiometry (MAICO ST 20, MAICO Diagnostic, GmBh, Berlin) confirmed normal hearing thresholds in all participants. Self-reported handedness was validated using a German translation of the Annett Handedness Questionnaire (Annett, 1970). To ensure group comparability with regard to general cognitive abilities, intelligence was evaluated using the Mehrfachwahl-Wortschatz-Intelligenztest (MWT-B; Lehrl, 2005). Musical aptitude was estimated using the Advanced Measures of Music Audiation (AMMA; Gordon 1989). The AMMA consists of 30 pairs of piano melodies. Participants are asked to decide whether the two melodies are identical, different in rhythmical patterns, or different in tonal patterns. The test results in a rhythmical score, a tonal score, and a total score (which equals the sum of rhythmical and tonal score). Participants’ characteristics of the two groups are given in Table 1.

**Table 1.** Participant Characteristics

|  | Absolute Pitch Musicians<br>(*n* = 54) | Non-Absolute Pitch Musicians<br>(*n* = 51) |
| --- | --- | --- |
| Sex |  |  |
| Female | 27 | 24 |
| Male | 27 | 27 |
| Age (years) | 26.67 (5.49) | 25.37 (4.49) |
| Handedness |  |  |
| Right-handed | 47 | 46 |
| Left-handed | 4 | 4 |
| Both-handed | 3 | 1 |
| Intelligence (MWT-B) <sup>a</sup> | 27.69 (5.10) | 29.10 (4.64) |
| Age of Onset of Musical Training (years) | 5.93 (2.39) | 6.49 (2.44) |
| Lifetime Cumulative Training (hours) <sup>b</sup> | 1.66 (1.22) | 1.35 (0.96) |
| Musical Aptitude (AMMA) <sup>a</sup> – total | 66.11 (6.31) | 63.35 (6.86) |
| Musical Aptitude (AMMA) <sup>a</sup> – tonal | 32.33 (3.75) | 30.45 (4.13) |
| Musical Aptitude (AMMA) <sup>a</sup> –<br>rhythmical | 33.78 (2.83) | 32.90 (3.03) |
| Pitch-labeling Task (%) | 76.41 (19.55) | 24.04 (18.92) |

*Annotations.* Continuous measures are given as mean (standard deviations in parentheses). Abbreviations: MWT-B = Mehrfachwahl-Wortschatz-Intelligenztest, AMMA = Advanced Measures of Music Audiation.
<sup>a</sup> Raw scores
<sup>b</sup> Units are given in 10,000

The study was approved by the ethics committee of the canton of Zurich (http://www.kek.zh.ch) and was performed in accordance with the Declaration of Helsinki. Written informed consent was obtained from all participants.

### Pitch-Labeling Task

Pitch-labeling ability was evaluated using a web-based adaptation of a task previously applied by our research group (Oechslin, Meyer, *et al*., 2010). The task consisted of 108 trials with pure tones ranging from C3 to B5 (tuning: A4 = 440 Hz). In every trial, 2,000 ms of Brownian noise, a 500-ms pure tone, and again 2,000 ms of Brownian noise were sequentially presented. Overall, each tone appeared three times in a pseudorandomized presentation order: No tone was repeated in successive trials.

Participants were asked to identify the pitch class (chroma, e.g., G) and octave (e.g., 4) of the pure tone by choosing one label (e.g., G4) out of a list of all possible labels (C3 to B5). Trials could be terminated by clicking on a button and had a maximal duration of 15,000 ms. Pitch-labeling accuracy was calculated as the percentage of correctly identified pitch classes. Octave errors were not penalized, resulting in a chance level of 8.3 %.

### EEG Recording and Preprocessing

For EEG recording, participants were seated in an electrically shielded, dimly lit room and instructed to relax with their eyes closed. The eyes-closed resting-state EEG was recorded for three minutes with a sampling rate of 1,000 Hz using a BrainAmp amplifier (Brainproducts, Munich, Germany). The 32 silver/silver-chloride electrodes were mounted on an electrode cap (Easycap, Herrsching, Germany) according to a subset of the 10/10 system (Fp1, Fp2, F7, F3, Fz, F4, F8, FT7, FC3, FCz, FC4, FT8, T7, C3, Cz, C4, T8, TP9, TP7, CP3, CPz, CP4, TP8, TP10, P7, P3, Pz, P4, P8, O1, Oz, and O2). An electrode on the tip of the nose served both as an online and offline reference. During EEG acquisition, a bandpass filter of 0.1 – 100 Hz was applied, and electrode impedances were kept below 10 kΩ by application of an abrasive and electrically conductive gel. After recording of the resting-state EEG, participants performed a passive auditory oddball task (for details see: Greber *et al*., 2018) and a pitch processing task (for details see: Leipold, Brauchli, Greber, & Jäncke, 2019; Leipold, Greber, Sele, *et al*., 2019).

The acquired EEG data were preprocessed using the BrainVision Analyzer software package (Version 2.1, https://www.brainproducts.com/). First, a bandpass filter between 1 and 20 Hz (48 dB/octave), and a notch filter of 50 Hz were applied. Then, a restricted infomax independent component analysis (ICA; (Jung *et al*., 2000)) was used to correct eye movement artifacts. Noisy channels were excluded from the ICA and interpolated after ICA correction. Finally, the continuous EEG was divided into segments of 2,000 ms. Segments with a voltage gradient > 100 μV/ms, an amplitude > 200 μV, or an amplitude < −200 μV were automatically rejected, resulting in a minimum of 62 and maximum of 90 artifact-free segments per participant.

### EEG Source-Level Estimation

To compute source-level EEG functional connectivity, the EEG segments were imported into the sLORETA/eLORETA (standardized/exact low-resolution brain electromagnetic tomography) toolbox (Version v20151222, http://www.uzh.ch/keyinst/loreta.htm). There, the neural generators of the electric potential differences on the scalp were estimated using the eLORETA algorithm (Pascual-Marqui *et al*., 2011), a linear, weighted minimum inverse solution with exact localization to point sources. eLORETA uses a realistically shaped head model (Fuchs *et al*., 2002) based on the Montreal Neurological Institute (MNI) 152 template (Mazziotta *et al*., 2001) for source reconstruction. The three-dimensional cortical solution space is restricted to gray matter and comprises 6239 voxels with a size of 5 x 5 x 5 mm^3^. To validate the accuracy of the source reconstruction, we used EEG data from the auditory oddball task performed by the same participants immediately after the resting-state recording (Greber *et al*., 2018). Based on the grand average over all participants, we estimated the source activity of the N1 component of the event-related potential evoked by the standard tone C4 (piano tone with a fundamental frequency f_0_ = 264 Hz). Maximal current density was localized in the transverse temporal gyrus (xyz coordinates: 65, −15, 10; Brodmann Area [BA] 42), confirming that the eLORETA algorithm performed as intended on our data.

### EEG Source-Level Connectivity

Based on the estimated source-level activity of the EEG resting-state segments, we conducted two types of connectivity analyses: an ROI-based replication analysis and an exploratory whole-brain network analysis. For both analyses, source-level EEG functional connectivity was evaluated with lagged phase synchronization, lagged linear connectivity, and instantaneous linear connectivity. Lagged phase synchronization is the connectivity measure used in Elmer *et al*.’s (2015) study. It quantifies the similarity between the normalized Fourier transforms of two signals (i.e. the time series in one brain region and the time series in another brain region) at a specific frequency after removal of the instantaneous, zero-phase contribution. It is a measure of non-linear dependency, is insensitive to amplitude information, and takes values between zero (independence) and one (perfect similarity). The two additionally analyzed connectivity measures, on the other hand, describe the linear coherence-type similarity between two signals at a specific frequency and incorporate both phase and amplitude information. They are also non-negative but have no upper bound (i.e., infinity corresponds to perfect similarity). Their sum equals the total linear connectivity, whereby the lagged part is only minimally affected by non-physiological artifacts, as for example volume conduction and the low spatial resolution (Pascual-Marqui, 2007; Pascual-Marqui *et al*., 2011). Contrary to lagged measures, instantaneous measures of connectivity are contaminated with non-physiological artifacts. Yet, they have been shown to surpass lagged measures in biometric identification of individuals (Valizadeh *et al*., 2019) and in the proportion of variance explained by structural connectivity (Finger *et al*., 2016). Furthermore, instantaneous connectivity measures have been successfully used to obtain meaningful expertise-related resting-state networks in previous studies (e.g., Jäncke & Langer, 2011; Klein *et al*., 2015, 2018). Hence, (near) zero-lag dependency seems to carry some relevant physiological information that is not fully captured by lagged measures. The use of the described connectivity measures enabled us to examine phase-only and phase-amplitude, as well as zero-lag and lagged connectivity differences between the two groups.

For the replication analysis, we defined four ROIs in the cortical solution space using the centroid voxels reported in Elmer *et al*.’s study (2015; see Figure 2-A). In each hemisphere, one ROI was placed in the auditory cortex (BA 41/42, xyz coordinates: ±54, −25, 10) and one ROI was placed in the DLPFC (BA 9/10/46, xyz coordinates: ±25, 45, 24). As in the original study, EEG functional connectivity between the two ROIs in each hemisphere was evaluated in the theta frequency band (4 – 7 Hz).

For the exploratory whole-brain network analysis, we computed lagged phase synchronization, linear lagged connectivity, and linear instantaneous connectivity between the centroid voxels of all 84 BAs as implemented in the sLoreta/eLoreta toolbox. Here, we included four frequency bands: theta (4 – 7 Hz), alpha (8 – 12 Hz), lower beta (13 – 21 Hz), and upper beta (22-30 Hz).

### Data Availability

Demographic and behavioral data, EEG raw data, EEG connectivity values, and mean network values are available online at https://osf.io/hbz28/?view_only=53d60b0dc7844179a4b15526d7736bca.

### Statistical Analysis

We performed (1) statistical analyses of the demographic and behavioral data, (2) replication analyses of the EEG functional connectivity between the auditory cortex and the DLPFC, and (3) network-based analyses of whole-brain EEG functional connectivity.

If not otherwise specified, the analyses were performed using R (version 3.4.3; https://www.r-project.org; R Core Team, 2017). Frequentist Analyses of variance (ANOVAs) were computed using the R package ez (version 4.4.0; https://cran.r-project.org/web/packages/ez/index.html; Lawrence, 2016). Unless otherwise stated, the significance level α was set to 0.05. We report effect sizes as generalized eta-squared η^2^_G_ (Bakeman, 2005) for ANOVAs and as Cohen’s *d* (Cohen, 1988) for *t*-tests.

#### Statistical Analyses of Demographic and Behavioral Data

The musical aptitude test AMMA was analyzed with a 2 x 2 ANOVA with factors Group (AP and non-AP) and Score Subtype (tonal and rhythmical). All other participant characteristics and behavioral data were analyzed using two-tailed Welch’s *t*-tests.

#### EEG ROI-Based Replication Analyses

For the ROI-based replication analysis, we used both frequentist and Bayesian statistics. The frequentist analysis exactly replicated the statistical methods used in the original study (Elmer *et al*., 2015). However, frequentist analyses are limited in that they only permit the rejection of the null hypothesis (H0) but not of the alternative hypothesis (H1). Non-significant results cannot be interpreted as evidence for the absence of an effect. In contrast, Bayesian statistics quantify the evidence both for and against H0 (Rouder *et al*., 2009; Dienes, 2011, 2014; Lee & Wagenmakers, 2013), which is especially useful for the interpretation of non-significant results (Dienes, 2014) and for the evaluation of replication success (Anderson & Maxwell, 2016). Thus, we computed Bayes factors in addition to the frequentist analysis. Bayes factors compare the (marginal) likelihood of the data under one hypothesis (e.g., H0) with the (marginal) likelihood of the data under another hypothesis (e.g., H1). The relative evidence for one hypothesis as expressed by a Bayes factor can be readily interpreted: A Bayes factor of BF_10_ = 5 (or the inverse 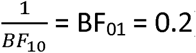) means that the data is five times more likely to occur under H1 than under H0.

For the frequentist replication analyses, the lagged phase synchronization values were subjected to a 2 x 2 ANOVA with factors Group (AP and non-AP) and Hemisphere (left and right). We also computed a one-tailed Welch’s *t*-test to specifically examine the group difference in the left hemisphere, in which the original study found higher connectivity in AP. In addition to the group statistics, we computed one-sided partial correlations for AP musicians between pitch-labeling accuracy and left hemispheric connectivity adjusted for the age of onset of musical training.

Bayes factors for Bayesian ANOVAs (BANOVAs), Bayesian *t*-tests, and Bayesian correlations were computed using the R package BayesFactor (version 0.9.12-4.2; https://cran.r-project.org/web/packages/BayesFactor/index.html; Morey *et al*., 2018). We used the default priors (a Cauchy distribution centered around zero with a scale parameter of 0.707) with the default number of iterations (n = 10,000). Since effect sizes are often inflated in studies with small sample sizes, we refrained from using scale-informed priors based on the effect sizes of the original study (Ioannidis, 2008; Button *et al*., 2013; Halsey *et al*., 2015). To confirm the robustness of the results, we tested a variety of additional priors with scale parameters between 0.5 (medium) and 1 (ultrawide); the results suggested the same conclusions as reported here. For the BANOVAs, Bayes factors of the two main effects (group and hemisphere) were assessed by comparing the model with one factor (e.g., group + subject) to the model with both factors (e.g., group + hemisphere + subject). Interaction effects were assessed by comparing the full model (group + hemisphere + group * deviation + subject) to the model without the interaction effect (group + hemisphere + subject). The Bayes factors reported for the one-sided correlation analyses do not account for the age of onset of musical training.

Extending the analyses of the original study, we analyzed two additional connectivity measures (lagged linear connectivity and instantaneous linear connectivity) to check whether the effect generalizes to other measures of functional connectivity. We report one-sided Welch’s *t*-tests and Bayesian *t*-tests, and correlations for both hemispheres for all measures as described above.

#### EEG Whole-Brain Network-Based Analyses

While the literature- and hypothesis-based definition of two centroids per hemisphere mitigates the multiple comparison problem, it carries the risk of missing other meaningful connections. For this reason, we additionally applied a less-restrictive, exploratory approach to investigate the resting-state EEG data. The whole-brain eLORETA output matrices (84 centroids of 84 BAs) were subjected to group comparisons using the network-based statistic (NBS) toolbox (Zalesky *et al*., 2010; http://www.nitrc.org/projects/nbs/) in MATLAB (version R2017b; https://www.mathworks.com/products/matlab.html). The analysis was performed separately for the four frequency bands of interest (theta, alpha, lower beta, and upper beta) and the three connectivity measures (lagged phase synchronization, lagged linear connectivity, and instantaneous linear connectivity). The NBS method provides a control for the family-wise error (FWE) rate when testing each connection between many ROIs. It applies the same principles as nonparametric cluster-based thresholding conventionally used in fMRI analyses (Nichols & Holmes, 2001). By considering interconnectedness in the topological space, NBS treats networks holistically and does not declare significance for individual connections.

To compare the individual connectivity matrices between the groups, we used the *t*-test module in NBS for both one-tailed contrasts (1, −1 and −1, 1). First, this module computed *t*-test statistics for each pairwise association between the 84 ROIs. Edges exceeding a specified threshold formed a suprathreshold network if connected with each other. The size (i.e., the number of edges) of the largest observed suprathreshold network was subjected to permutation testing. For a total of 5,000 permutations, the group labels of the participants were randomly exchanged, and the analysis was repeated using the same threshold. From each permutation step, the size of the largest suprathreshold network was stored to form an empirical estimate of the null distribution. The p-value of the observed network was estimated by counting the permutations that yielded the same or a bigger maximal network size and dividing this count by the total number of permutations. Thus, the reported p-values are FWE corrected only for the number of ROIs. We applied no additional correction for the number of NBS tests performed because of the exploratory nature of the analysis.

Because we were interested in middle (d ≈ 0.4) to large (*d* ≈ 0.8) effect sizes on the level of individual links, we tested the connectivity matrices for the corresponding thresholds between *t* = 2.0 and *t* = 4.0 in increments of 0.1. For each separate analysis (four frequency bands, three connectivity measures, two contrasts), we report all thresholds at which a statistically significant network (*p* <. 05) emerged and describe the network at one of these thresholds in detail. This threshold was chosen by inspection of the networks: The number of nodes should neither be too great nor too small for interpretation, and the network should not have disintegrated into multiple subnetworks. The reported significant networks were visualized using the BrainNet Viewer software (version 1.53; http://www.nitrc.org/projects/bnv/) in MATLAB (Version R2017b, https://www.mathworks.com/products/matlab.html). The Harvard-Oxford cortical atlas and the Juelich Histological atlas as implemented in FSL (http://fsl.fmrib.ox.ac.uk/fsl/fslwiki/FSL) were used to specify the brain regions underlying the involved nodes.

After identification of the networks, we analyzed the relationship between the corresponding mean network values and pitch-labeling performance using R (version 3.4.3; https://www.r-project.org; R Core Team, 2017). We computed both frequentist and Bayesian correlations (two-sided, non-partial) for each group separately.

## Results

### Results of Demographic and Behavioral Analyses

AP and non-AP musicians were comparable in age (*t*_(100.97)_ = 1.33, *p* =. 19, *d* = 0.26), intelligence (*t*_(102.86)_ = −1.49, *p* =. 14, *d* = 0.29), age of onset of musical training (*t*_(102.42)_ = −1.20, *p* =. 23, *d* = 0.23), and cumulative musical training hours over the lifespan (*t*_(99.71)_ = 1.43, *p* =. 16, *d* = 0.28). The analysis of the AMMA scores (measuring musical aptitude) yielded a main effect of Group (*F*_(1, 103)_ = 4.60, *p* =. 034, η^2^_G_ = 0.04), a main effect of Score Subtype (*F*_(1, 103)_ = 79.27, *p* <. 001, η^2^_G_ = 0.07), and an interaction effect (*F*_(1, 103)_ = 5.37, *p* =. 023, η^2^_G_ = 0.005). Post hoc *t*-tests (Bonferroni corrected α = 0.25) revealed that the AP musicians were comparable to non-AP musicians in the rhythmical score (*t*_(101.38)_ = 1.53, *p* =. 13, *d* = 0.30) but had a higher tonal score (*t*_(100.61)_ = 2.44, *p* =. 016, *d* = 0.48). As expected, AP musicians outperformed non-AP musicians in the pitch-labeling task (*t*_(102.93)_ = 13.95, *p* <. 001, *d* = 2.72; see Figure 1).

**Figure 1.**
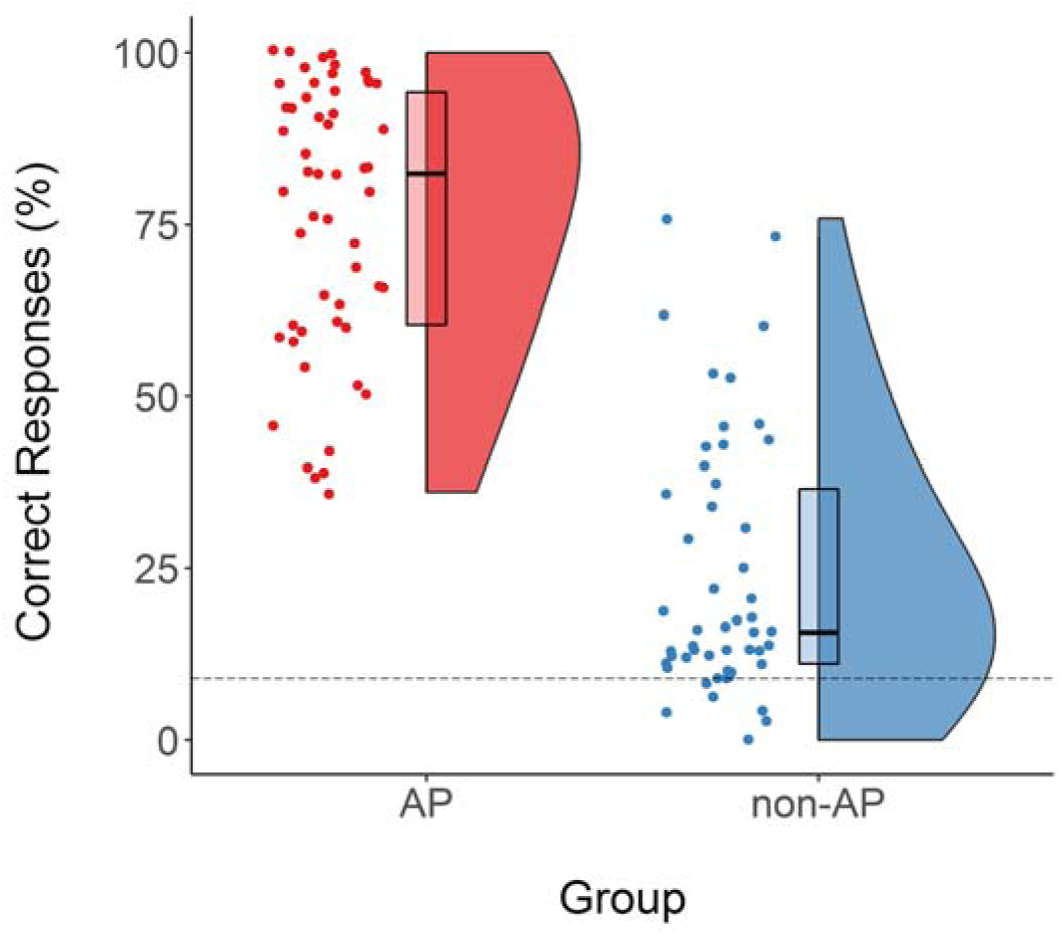
Pitch-labeling scores for AP (n = 54) and non-AP (n = 51) musicians. Because octave errors were disregarded, the chance level was 8.33 % (dashed line). Abbreviations: AP = absolute pitch.

### Results of ROI-Based Replication Analyses

The ROI-based replication analysis of lagged phase synchronization – the measure used in the original study (Elmer *et al*., 2015) – in the theta frequency band between the auditory cortex and the DLPFC revealed no evidence for a main effect of Group (*F*_(1, 103)_ = 1.86, *p* =. 18, η^2^_G_ = 0.01, BF_01_ = 3.39), no evidence for a main effect of Hemisphere (*F*_(1, 103)_ = 0.06, *p* =. 81, η^2^_G_ < 0.001, BF_01_= 8.90), and no evidence for a Group x Hemisphere interaction (*F*_(1, 103)_ = 0.01, *p* =. 91, η^2^_G_ < 0.001, BF_01_ = 6.69). Lagged-synchronization values are shown in Figure 2-B and posterior distributions of the BANOVA are illustrated in Figure 2-D. The planned one-tailed *t*-test did not reveal evidence for a difference between the two groups in the left hemisphere (*t*_(102.75)_ = −0.90, *p* =. 81, *d* = 0.18, BF_01_ = 8.49; see Figure 2-B). There was also no evidence for a positive relationship between pitch-labeling performance and left-hemispheric lagged phase synchronization in AP musicians (*r*_*p*_ = -.034, *p* = 1.00, BF_01_ = 4.26; see Figure 2-C).

**Figure 2.**
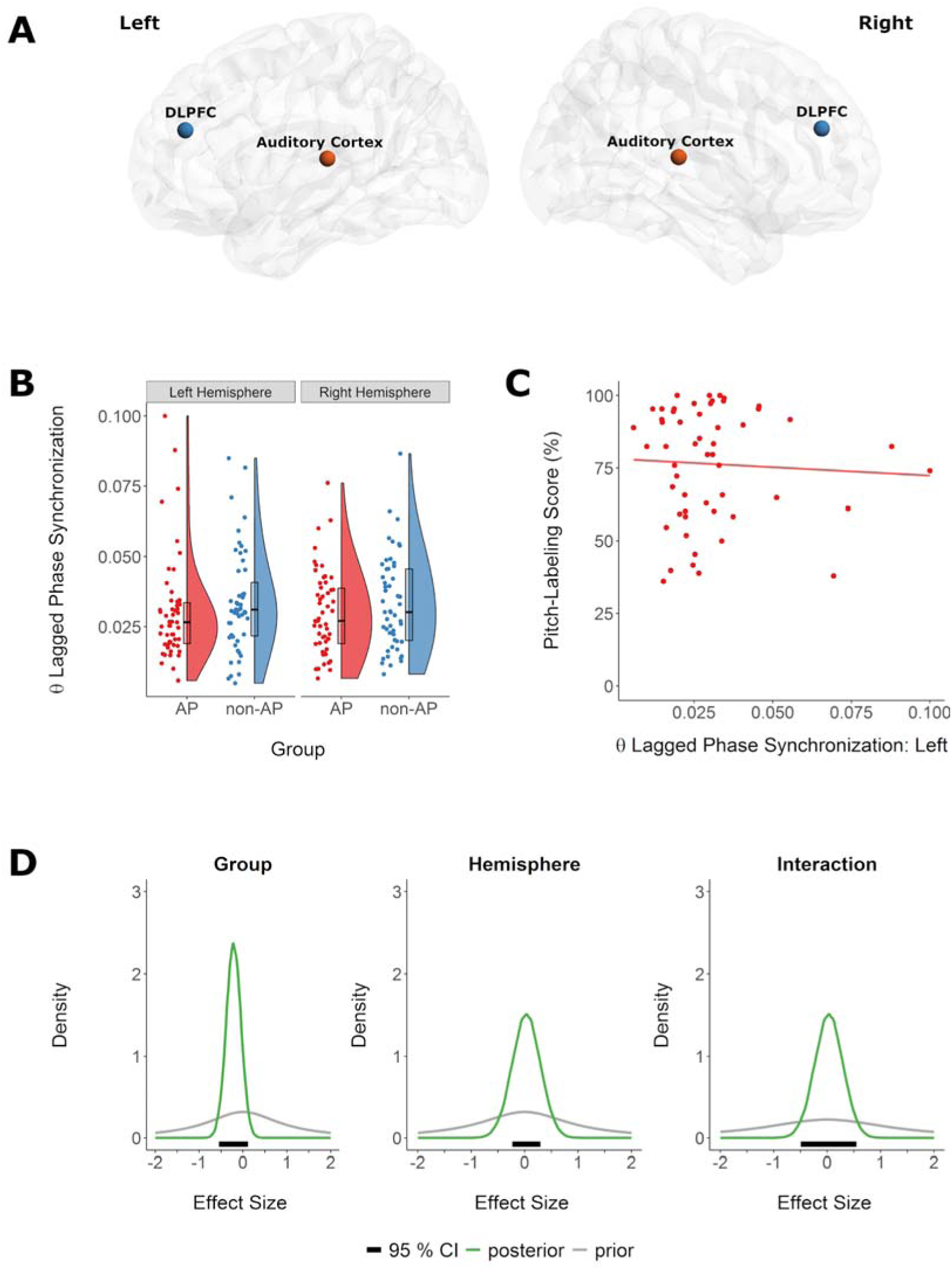
Replication analysis of theta lagged phase synchronization between the auditory cortex and the dorsolateral prefrontal cortex (DLPFC) during EEG resting state. A) Localization of the four regions of interests (ROIs). B) There was no evidence for a difference in theta lagged phase synchronization values between absolute pitch musicians (red) and non-AP musicians (blue). C) There was no evidence for a positive correlation between left-hemispheric theta lagged phase synchronization and performance in the pitch-labeling task in AP musicians. D) Prior (gray) and posterior (green) distributions of the standardized effects (relative to the standard deviation of the error term) of the factors Group and Hemisphere on theta lagged phase synchronization. The Bayesian 95 % credible interval (CI) describes the interval that includes the true value with a probability of 95 %, given the data and the assumed model.

Additional analyses of resting-state connectivity between the auditory cortex and the DLPFC in AP musicians based on lagged linear connectivity and instantaneous linear connectivity also revealed no evidence for differences between the two groups. All results of the group comparisons for each hemisphere are shown in detail in Table 2. There was also no evidence for a positive relationship between pitch-labeling performance and resting-state connectivity between the auditory cortex and the DLPFC in AP musicians. The results of the correlational analyses are shown in Table 3.

**Table 2.** Group Comparisons of Theta-Band Connectivity between the Auditory Cortex and the DLPFC.

| Connectivity Measure | Hemisphere | AP | Non-AP | *p*-value | Cohen's *d* | BF<sub>01</sub> |
| --- | --- | --- | --- | --- | --- | --- |
| Lagged phase synchronization | Right | 0.030<br>(0.015) | 0.033<br>(0.017) | .88 | 0.23 | 9.77 |
|  | Left | 0.030<br>(0.018) | 0.034<br>(0.018) | .81 | 0.18 | 8.49 |
| Lagged linear connectivity | Right | 0.046<br>(0.029) | 0.049<br>(0.031) | .70 | 0.11 | 6.96 |
|  | Left | 0.043<br>(0.028) | 0.051<br>(0.032) | .91 | 0.27 | 10.64 |
| Instantaneous linear connectivity | Right | 0.967<br>(0.773) | 0.810<br>(0.466) | .10 | 0.25 | 1.37 |
|  | Left | 0.787<br>(0.415) | 0.756<br>(0.433) | .35 | 0.07 | 3.55 |

*Annotations.* One-sided Welch's *t*-tests were applied to compare AP and non-AP musicians (hypothesis AP > non-AP). Group values for AP and non-AP musicians are given as mean (standard deviation in parentheses). Lagged phase synchronization was the measure used in the original study (Elmer et al., 2015). Abbreviations: AP = absolute pitch, DLPFC = dorsolateral prefrontal cortex.

**Table 3.** Correlations between Pitch-Labeling Performance and Theta-Band Connectivity in AP Musicians.

| Connectivity Measure | Hemisphere | $r$ | $p_{one-sided}$ | BF<sub>01</sub> |
| --- | --- | --- | --- | --- |
| Lagged phase synchronization | Right | 0.160 | .12 | 0.99 |
|  | Left | -0.034 | 1.00 | 4.26 |
| Lagged linear connectivity | Right | 0.001 | 0.50 | 2.49 |
|  | Left | -0.157 | 1.00 | 3.86 |
| Instantaneous linear connectivity | Right | -0.199 | 1.00 | 6.70 |
|  | Left | -0.054 | 1.00 | 3.86 |

*Annotations.* One-sided partial correlations ( $r$ and $p$ adjusted for age of onset of musical training; hypothesis higher pitch-labeling score is associated with stronger connectivity). Theta-band connectivity was evaluated between the auditory cortex and the DLPFC. Lagged phase synchronization was the measure used in the original study (Elmer et al., 2015). Abbreviations: AP = absolute pitch, DLPFC = dorsolateral prefrontal cortex.

### Results of Whole-Brain Network-Based Analyses

The network-based analyses of the 84-ROI connectivity matrices revealed group differences in three measure x frequency combinations (see Figure 3): AP musicians showed hyperconnected resting-state networks in (A) lagged linear connectivity in lower beta, (B) in instantaneous linear connectivity in lower beta, and (C) in instantaneous linear connectivity in theta. No significant networks were observed in lagged phase synchronization or in any of the other tested frequency bands of lagged and instantaneous linear connectivity. The analyses did also not reveal any networks with decreased connectivity in AP musicians compared to non-AP musicians.

**Figure 3.**
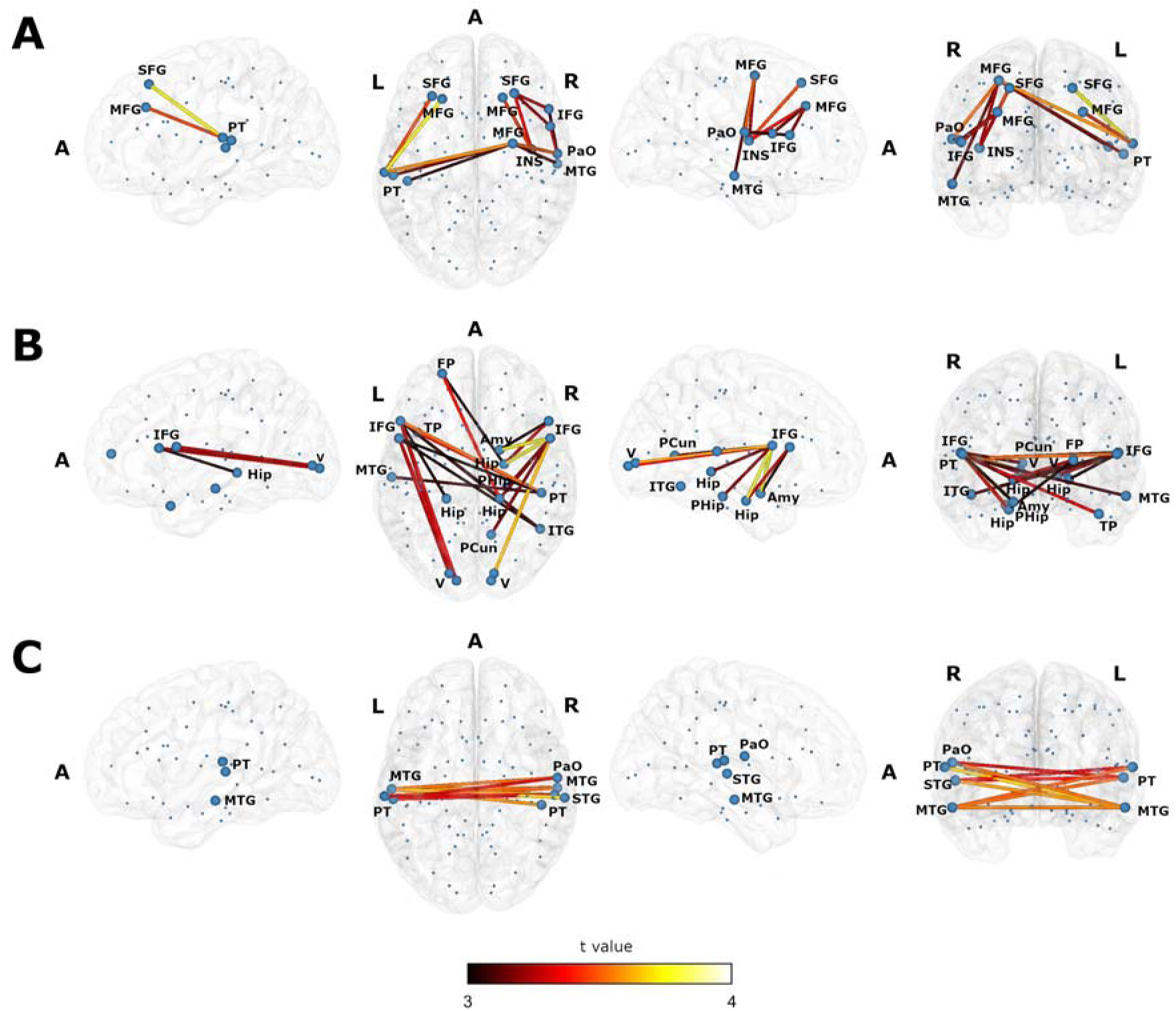
Lateral, axial and coronal views of the three obtained resting-state networks. All three networks show increased undirected connectivity in AP musicians compared to non-AP musicians. Blue spheres represent the centroids of the 84 Brodmann Areas. Nodes contributing to the network are depicted by enlarged spheres. The color of the edges corresponds to the *t*-value. A) The network in lagged linear connectivity in lower beta. B) The network in instantaneous linear connectivity in lower beta. C) The network in instantaneous linear connectivity in theta. Abbreviations: A = anterior, Amy = Amygdala, FP = frontal pole, Hip = hippocampus subiculum, IFG = inferior frontal gyrus, INS = insular cortex, ITG = inferior temporal gyrus, L = left hemisphere, MFG = middle frontal gyrus, PaO = parietal operculum, PCun = precuneus cortex, PHip = parahippocampal gyrus, PT = planum temporale, R = right hemisphere, SFG = superior frontal gyrus, STG = superior temporal gyrus, TP = temporal pole, V = visual cortex.

A. In lagged linear connectivity in the lower beta frequency band, statistically significant networks were found for all tested thresholds between *t* = 2.0 (76 nodes, 423 edges) and *t* = 3.7 (2 nodes, 1 edge). We report the network at *t* = 3.0, visualized in Figure 3-A. At this threshold, 13 nodes and 14 edges contributed to the network (*p* =. 037, FWE corrected for the number of ROIs). The brain regions underlying the involved nodes are listed in Table 4. Nodes in the left temporal lobe (auditory regions, planum temporale) were connected to nodes in the frontal lobe both intrahemispherically (left middle and superior frontal gyrus) and interhemispherically (right middle/ superior frontal gyrus, BA 6). Within the right hemisphere, nodes in the frontal lobe (middle frontal gyrus, superior frontal gyrus, inferior frontal gyrus), in the parietal operculum, in the insular cortex, and in the middle temporal gyrus contributed to the network. Two-sided correlations revealed no evidence for a relationship between mean network values and pitch-labeling performance within AP musicians (*r* =. 095, *p* =. 49, BF_01_ = 2.63) or within non-AP musicians (*r* =. 075, *p* =. 60, BF_01_ = 2.80).
B. In instantaneous linear connectivity in the lower beta frequency band, significant networks were obtained at thresholds between *t* = 2.0 (77 nodes, 411 edges) and *t* = 3.0 (19 nodes, 23 edges), and at *t* = 3.6 (4 nodes, 3 edges) and *t* = 3.7 (3 nodes, 2 edges). The relatively widespread network at *t* = 3.0 (*p* =. 044, FWE corrected for number of ROIs; see Table 5 and Figure 3 B) consisted of nodes in the occipital lobe (visual cortex, occipital pole, precuneus), in subcortical regions (hippocampal and parahippocampal regions), in the temporal lobe (inferior temporal gyrus, middle temporal gyrus, temporal pole, planum temporale/auditory cortex), and in the frontal lobe (frontal pole, inferior frontal gyrus). There was no evidence for a correlation between mean network values and pitch-labeling performance within the AP group (*r* =. 004, *p* =. 97, BF_01_ = 3.25) or within the non-AP group (*r* =. 008, *p* =. 96, BF_01_ = 3.17).
C. In instantaneous linear connectivity in the theta frequency band, NBS revealed significant networks at thresholds between *t* = 3.1 (11 nodes, 15 edges) and *t* = 3.5 (7 nodes, 6 edges). At a middle-level threshold of *t* = 3.3, the network (*p* =. 032, FWE corrected for number of ROIs) compromised of 8 nodes in temporal and perisylvian regions (middle temporal gyrus, superior temporal gyrus, planum temporale, auditory cortex, and parietal operculum) and of 10 interhemispheric connections. The network nodes are described in detail in Table 6, and the network is visualized in Figure 3-C. Similar to the other two networks, there was no evidence for a relationship between mean network values and pitch-labeling performance in either AP (*r* =. 070, *p* =. 63, BF_01_ = 2.92) or non-AP musicians (*r* = -.16, *p* =. 27, BF_01_ = 1.83).

**Table 4.** Brain regions underlying the centroid voxel coordinates of the BAs constituting the lower-beta linear lagged connectivity network (*t* threshold = 3.0 associated with a Cohen’s *d* = 0.59).

| MNI coordinates (x, y, z) | Brain region | Brodmann Area |
| --- | --- | --- |
| -22, 28, 49 | left superior frontal gyrus | BA 8 |
| -29, 30, 33 | left middle frontal gyrus | BA 9 |
| -56, -25, 5 | left planum temporale/ primary auditory cortex | BA 41 |
| -46, -29, 10 | left planum temporale/ primary auditory cortex | BA 41 |
| -62, -23, 12 | left planum temporale | BA 42 |
| 27, -3, 54 | right middle frontal gyrus/ superior frontal gyrus | BA 6 |
| 20, 29, 49 | right superior frontal gyrus | BA 8 |
| 28, 32, 33 | right middle frontal gyrus | BA 9 |
| 40, -7, 9 | right insular cortex | BA 13 |
| 58, -17, -15 | right middle temporal gyrus | BA 21 |
| 58, -10, 15 | right secondary somatosensory cortex/ parietal operculum | BA 42 |
| 53, 9, 14 | right inferior frontal gyrus (Broca's area BA44) | BA 44 |
| 52, 21, 13 | right inferior frontal gyrus (Broca's area BA44/BA45) | BA 44/45 |

*Annotations.* Nodes were assigned to brain regions based on the Harvard-Oxford cortical atlas and the Juelich Histological atlas. Brodmann areas (BAs) refer to the LORETA output.

**Table 5.** Brain regions underlying the centroid voxel coordinates of the BAs constituting the lower-beta linear instantaneous connectivity network (*t* threshold = 3.0 associated with a Cohen’s *d* = 0.59).

| MNI coordinates (x, y, z) | Brain region | Brodmann Area |
| --- | --- | --- |
| -22, 54, 9 | left frontal pole | BA 10 |
| -12, -90, -1 | left visual cortex (V1, V2, V3)/occipital pole | BA 17 |
| -17, -85, 1 | left visual cortex (V3) | BA 17 |
| -57, -18, -15 | left middle temporal gyrus | BA 21 |
| -19, -33, -4 | left hippocampus subiculum | BA 27 |
| -39, 13, -27 | left temporal pole | BA 38 |
| -52, 9, 14 | left inferior frontal gyrus (Broca's area<br>BA44) | BA 44 |
| -51, 21, 13 | left inferior frontal gyrus (Broca's area<br>BA44) | BA 44/45 |
| 12, -90, 0 | right visual cortex (V1)/occipital pole | BA 17 |
| 14, -85, 2 | right visual cortex (V1) | BA 17 |
| 18, -33, -4 | right hippocampus subiculum | BA 27 |
| 21, -9, -24 | right hippocampus subiculum | BA 28 |
| 12, -58, 7 | right visual cortex (V1)/ precuneus cortex | BA 30 |
| 18, 1, -19 | right amygdala superficial group/<br>parahippocampal gyrus | BA 34 |
| 23, -25, -21 | right parahippocampal gyrus | BA 35 |
| 46, -54, -14 | right inferior temporal gyrus/ temporal<br>occipital fusiform cortex | BA 37 |
| 47, -29, 10 | right planum temporale/ primary auditory<br>cortex | BA 41 |
| 53, 9, 14 | right inferior frontal gyrus (Broca's area<br>BA44) | BA 44 |
| 52, 21, 13 | right inferior frontal gyrus (Broca's area<br>BA44/BA45) | BA 44/45 |

*Annotations.* Nodes were assigned to brain regions based on the Harvard-Oxford cortical atlas and the Juelich Histological atlas. Brodmann areas (BAs) refer to the LORETA output.

**Table 6.**
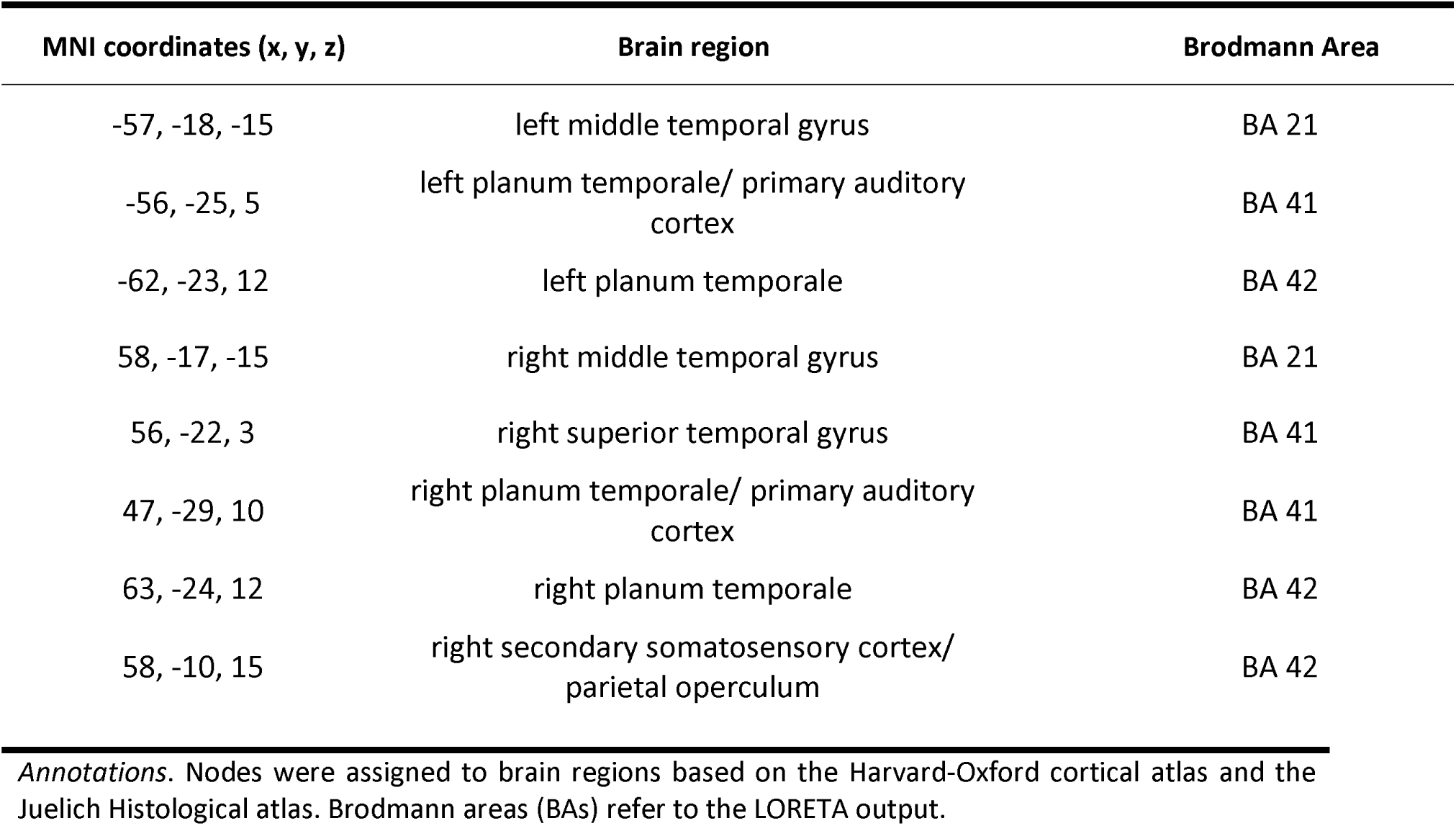
Brain regions underlying the centroid voxel coordinates of the BAs constituting the theta linear nstantaneous connectivity network (*t* threshold = 3.3 associated with a Cohen’s *d* = 0.65).

| MNI coordinates (x, y, z) | Brain region | Brodmann Area |
| --- | --- | --- |
| -57, -18, -15 | left middle temporal gyrus | BA 21 |
| -56, -25, 5 | left planum temporale/ primary auditory cortex | BA 41 |
| -62, -23, 12 | left planum temporale | BA 42 |
| 58, -17, -15 | right middle temporal gyrus | BA 21 |
| 56, -22, 3 | right superior temporal gyrus | BA 41 |
| 47, -29, 10 | right planum temporale/ primary auditory cortex | BA 41 |
| 63, -24, 12 | right planum temporale | BA 42 |
| 58, -10, 15 | right secondary somatosensory cortex/ parietal operculum | BA 42 |

*Annotations.* Nodes were assigned to brain regions based on the Harvard-Oxford cortical atlas and the Juelich Histological atlas. Brodmann areas (BAs) refer to the LORETA output.

## Discussion

This study investigated EEG resting-state connectivity in AP and non-AP musicians to provide insights into the role of perceptual and cognitive processes in AP. In a two-part analysis, we first attempted to replicate our previous finding of increased theta resting-state connectivity between the left auditory cortex and the left DLPFC (Elmer *et al*., 2015). In the second part, we performed an exploratory whole-brain analysis to evaluate whether the auditory cortex and the DLPFC are part of a larger AP-specific resting-state network.

In the ROI-based replication analysis, we found no evidence for an increase in theta-band lagged phase synchronization between the auditory cortex and the DLPFC in AP musicians compared to non-AP musicians. Bayes factor analyses favored the null hypothesis of no group differences (BF > 8). Similar results were obtained for two additionally analyzed connectivity measures. There was also no evidence for a positive relationship between pitch-labeling proficiency and left-hemispheric theta connectivity in the AP group. The whole-brain analysis provided some evidence in favor of hyperconnected networks in AP musicians in the theta and lower-beta frequency bands using instantaneous linear connectivity, and in the lower-beta frequency band using lagged linear connectivity.

### ROI-Based Replication Analyses: Auditory Cortex and DLPFC

In the ROI-based analysis, we did not replicate the previous finding (Elmer *et al*., 2015) of increased left-hemispheric temporo-frontal connectivity in AP musicians. This corresponds at least partly with previous reports on functional connectivity in AP. While the connectivity of the auditory cortex in AP has been addressed by several previous studies (Loui *et al*., 2011, 2012; e.g., Jäncke *et al*., 2012), much less is known about the DLPFC. For instance, a recent functional magnetic resonance imaging (fMRI) study found differential local connectivity patterns in the left auditory cortex during resting state but neither local nor global connectivity differences in the DLPFC between musicians with and without AP (Brauchli *et al*., 2019a). In another fMRI study, the Heschl’s gyrus was functionally connected to various auditory and non-auditory regions during passive tone listening in the AP group (Wengenroth *et al*., 2014). However, no evidence was found for an AP-specific synchronization between the auditory cortex and the DLPFC. Furthermore, Kim and Knösche (2017a) found no evidence for group differences in resting-state connectivity between the auditory cortex and seeds in the planum temporale, which is part of the dorsal auditory pathway between the auditory cortex and the DLPFC (Rauschecker & Scott, 2009). Alternatively, it has been proposed that the ventral pathway projecting to the inferior frontal gyrus via the anterior temporal lobe might play a more important role in AP processing than the DLPFC (Kim & Knösche, 2017a, 2017b; Leipold, Greber, & Elmer, 2019). The only other study besides Elmer *et al*. (2015) providing some evidence for the importance of a dorsal connection between auditory and frontal regions in AP found a leftward asymmetry of fractional anisotropy measures of the arcuate fasciculus in AP musicians but not in non-AP musicians or non-musicians (Oechslin, Imfeld, *et al*., 2010). The arcuate fasciculus structurally connects the posterior superior temporal gyrus and the prefrontal cortex (Makris *et al*., 2005). Taken together, there is not yet much support for increased connectivity between the auditory cortex and the DLPFC in AP, consistent with the results of the current study.

Some studies suggested that the DLPFC might be involved in the pitch-label association process in AP (Zatorre *et al*., 1998; Ohnishi *et al*., 2001; Bermudez & Zatorre, 2005; Levitin & Rogers, 2005). However, a recent fMRI study of our group did not observe an involvement of the DLPFC in AP during a pitch-processing task (Leipold, Brauchli, Greber, & Jäncke, 2019), casting doubt on the exact role of the DLPFC in pitch labeling. Activity in the DLPFC increased equally in musicians with and without AP between a listening and a labeling condition. Hence, we suggested that the activity in the DLPFC might actually reflect unspecific attentional or executive control processes rather than the label retrieval itself. The inconsistencies in DLPFC activation even during acoustic stimulation might explain to some extent why the increase in functional connectivity between the left auditory cortex and the left DLPFC could not be reliably detected during EEG resting state.

It is important to note that the DLPFC encompasses a rather large cortex region whose exact location and extension are not universally agreed upon (e.g., BA 8/9/46 O’Reilly, 2010; BA 8a/46 Rauschecker, 2011; BA 9/10/46 Teffer & Semendeferi, 2012; BA 9/46 Cieslik *et al*., 2013; BA 8/9/46 Plakke & Romanski, 2014). By considering only a single centroid within the DLPFC in our replication analysis, we cannot make statements about this broad region as a whole. We can only conclude that there was no evidence for an AP-specific increase in connectivity between the auditory cortex and the DLPFC as it was defined in the original study.

### Whole-Brain Network-Based Analyses

The exploratory whole-brain analyses yielded three resting-state networks with enhanced EEG connectivity (i.e., hyperconnectivity) in AP musicians compared to non-AP musicians. We did not find any evidence for networks with decreased connectivity in AP musicians. Several MRI studies have reported functional and structural hyperconnectivity in AP using a variety of both ROI-based and whole-brain methods (Loui *et al*., 2011, 2012; Wengenroth *et al*., 2014; Dohn *et al*., 2015; Kim & Knösche, 2017a; Brauchli *et al*., 2019a). On the other hand, there is also one report of reduced whole-brain structural connectivity (i.e., cortical thickness covariance) in AP musicians (Jäncke *et al*., 2012). Similarly, a recent EEG resting-state study observed global hypoconnectivity (i.e., lower clustering) in AP musicians on the electrode level (Wenhart *et al*., 2019). A recently published source-level EEG study, however, did not find any evidence for network differences between AP and non-AP musicians during resting state (Brauchli *et al*., 2019b). In contrast to our study, Brauchli and colleagues analyzed eyes-open instead of eyes-closed resting-state data. Taken together, there is little agreement on whether connectivity in AP musicians is increased, decreased or comparable to non-AP musicians. The greatly varying methods (e.g., imaging modality, structural vs functional, ROI-based vs whole-brain, electrode-level vs source-level, eyes-open vs eyes-closed, dependency measures, different types of connectivity and network analyses, different procedures for AP group assignment) may account for some of the diverging results. Resting-state connectivity of AP musicians might in particular be affected by the imaging modality. In addition to the inherent differences between fMRI and EEG regarding temporal and spatial resolution, there is no background noise during EEG recording. The fMRI scanner noise, on the other hand, might activate some pitch-labeling processes in AP musicians. Further research is necessary to disentangle hyper- and hypoconnectivity in AP and the influence of the respectively used methods.

The three networks we identified in our exploratory whole-brain analysis covered nodes in frontal, temporal, subcortical, and occipital brain regions. Common features across the three networks were the planum temporale, the inferior frontal gyrus, the parietal operculum, and the middle temporal gyrus. The planum temporale, a secondary auditory region posterior to the Heschl’s gyrus, has repeatedly been associated with AP (Schlaug *et al*., 1995; Zatorre *et al*., 1998; Keenan *et al*., 2001; Ohnishi *et al*., 2001; Luders *et al*., 2004; Wilson *et al*., 2009; Wengenroth *et al*., 2014; Leipold, Brauchli, Greber, & Jäncke, 2019; Burkhard *et al*., 2020). While its precise function in AP remains unknown, the planum temporale has been suspected to be involved in the matching of auditory input to internal templates (Griffiths & Warren, 2002). As recently put forward by Leipold, Brauchli, *et al*. (2019), a similar matching process specifically involving pitch templates might occur in the planum temporale of AP musicians during pitch labeling. The parietal operculum (secondary somatosensory cortex) has also been previously reported in connection with AP; its involvement was presumed to indicate sensorimotor integration (Wengenroth *et al*., 2014). However, considering the relatively low spatial resolution of EEG and the spatial closeness of the centroid voxels of the parietal operculum and the planum temporale, these nodes might not necessarily show selective neural activations of different brain regions in the present study. The inferior frontal gyrus has repeatedly been implicated in AP (Zatorre *et al*., 1998; Schulze *et al*., 2009; Wengenroth *et al*., 2014; Dohn *et al*., 2015; Leipold, Brauchli, Greber, & Jäncke, 2019; McKetton *et al*., 2019). Because its activity was either increased or decreased in AP musicians depending on the specific task, different functions have been attributed to it, such as a verbal component in AP processing (Wengenroth *et al*., 2014) or a working memory component in relative-pitch processing (Leipold, Brauchli, Greber, & Jäncke, 2019). Finally, the middle temporal gyrus has also been previously linked to AP (Zatorre *et al*., 1998; Loui *et al*., 2011; Wengenroth *et al*., 2014; Kim & Knösche, 2017a; Burkhard *et al*., 2019). The middle temporal gyrus participates in a multitude of functions (for an overview, consider Xu *et al*., 2015), including higher-order language processes (Friederici, 2002; Hickok & Poeppel, 2007; Oechslin, Meyer, *et al*., 2010). In the context of AP, the middle temporal gyrus has been proposed to play a role in accessing stored pitch templates (Loui *et al*., 2012), in categorizing perceived tones (Burkhard *et al*., 2019), or in recruiting multimodal codes for extracted pitch information (Zatorre *et al*., 1998).

The networks were found in the theta (4 – 7 Hz) and the lower-beta (13 Hz – 21 Hz) frequency range. A number of cognitive functions have been linked to these oscillation rhythms (for a review, see: Wang, 2010). For theta, these functions include working memory, memory encoding, and memory retrieval (Ward, 2003; Hsieh & Ranganath, 2014; Albouy *et al*., 2017), whereas the beta frequency band is involved in sensorimotor integration and top-down signaling (Engel & Fries, 2010; Siegel *et al*., 2012). These attributed functions are very well in accordance with the brain regions we found contributing to the AP-specific networks.

For all three networks, we found no evidence for a relationship between the mean network connectivity values and pitch-labeling scores within the group of AP musicians. Similarly, a recent fMRI resting-state study from our research project showed no significant correlations between the connectivity measures and pitch-labeling scores within the AP group (Brauchli *et al*., 2019). As argued there with reference to a large-scale behavioral study (Athos *et al*., 2007), a possible explanation for this lack of correlation might be that AP is a distinct rather than a continuous ability.

Overall, the nodes shared among the three networks corroborate the importance of perisylvian areas in AP, including prefrontal regions. The non-overlapping nodes of the networks might indicate the use of a widespread, possibly multisensory network. However, considering the number of exploratory NBS analyses that did not yield any evidence for group differences, the strength of evidence for hyperconnectivity during the eyes-closed resting-state remains weak.

### Limitations

Several general limitations apply to both the ROI-based and the whole-brain analysis. First, EEG source localization might be relatively imprecise when based on a low number of electrodes (Srinivasan *et al*., 1998; Baillet, 2017). To be sufficiently confident of the source reconstruction, we checked localization accuracy during acoustic stimulation, which confirmed that the eLORETA algorithm performed well on our data. Additionally, previous studies have verified the source reconstruction accuracy of the LORETA toolbox even for small numbers of scalp electrodes using intracranial electrode recording (Zumsteg *et al*., 2005, 2006). Second, the connectivity measures used in the analyses do not distinguish between direct and indirect connections (common input problem: Bastos & Schoffelen, 2016). Thus, connectivity between two nodes could have been mediated by a third source not included in the analysis. Lastly, caution must be applied when generalizing resting-state networks to active processing. As pointed out by Petersen and Sporns (2015), it could be that even brain networks activated by daily tasks (e.g., reading) are not necessarily expressed during resting state if, for instance, the contributing regions are also used by various other tasks. Thus, future connectivity analyses during active tasks are vital for a better understanding of the networks specifically involved in the process of pitch labeling in AP.

Additional limitations specifically apply to the ROI-based replication analysis. While both the current and the original study relied on self-reports with respect to group assignment to the AP and non-AP groups and retrospectively tested this group assignment using a pitch-labeling test, there are still some differences in terms of the used samples. First, we changed the assessment of the questionnaires and the pitch-labeling task from paper-pencil to online at home to lower the on-site testing workload for our participants. Second, due to the online implementation, the pitch-labeling task had to be slightly modified: Trials could last up to 15 s instead of a fixed duration of 5 s in the paper-pencil implementation of the original study. A pilot test showed that this modification was necessary for participants to be able to solve the multiple-choice format with 36 response options. Third, contrary to the original study, AP musicians scored higher than non-AP musicians in the tonal part of the musical aptitude test (AMMA) in the present study. Whilst statistically significant, this group difference was small in absolute numbers (less than 2 points out of a maximal score of 40 points), and the means were similar to those of the original sample. Finally, there was no overlap between the two groups in pitch-labeling scores in the original study (all non-AP musicians had less than 20 % correct, all AP-musicians had more than 35 % correct), but there was an overlap in our sample (highest score among non-AP musicians was 75.9 %, the lowest score among AP musicians was 36.1 %). This could be attributed to less homogenous groups but might also be due to the larger sample size or the longer trial duration in our pitch-labeling task: Because the participants had more time to respond, they might have used their relative-pitch ability to solve the task. It is conceivable that highly trained non-AP musicians can perform well under these circumstances. The difference between the two studies regarding the overlap in pitch-labeling scores seems to mostly stem from such well-performing non-AP musicians in the current study. AP musicians showed a similarly large range of pitch-labeling scores in the current and the original study. Furthermore, unpublished data from our lab suggests a strong correlation (r =0.7) between the online pitch-labeling test used in the current study and the on-site test used in the original study within AP musicians (n = 39). Although this correlation is strong there is still some unexplained variance, which might indicate that different cognitive functions have been involved during the performance of these different pitch-labeling task variants. Whether these suspected differences between the previous and the current study might be responsible for the different findings is disputable and should be examined in further experiments. We also found no evidence for a positive correlation between the pitch-labeling scores and the connectivity values of the ROI-based analysis within the AP group, which would have supported the importance of the connection between the auditory cortex and the DLPFC for AP.

### Conclusion

Using the ROIs defined in Elmer *et al*.’s (2015) study, we did not replicate an AP-specific increase in resting-state connectivity between the auditory cortex and the DLPFC in the theta frequency band. The exploratory whole-brain analyses provided some evidence for increased functional interactions among distributed brain areas in AP in the theta and lower-beta frequency bands. These areas comprised mainly of auditory and frontal brain regions but also included regions that engage in sensorimotor and visual processes. Future task-based studies using acoustic stimulation are necessary to confirm the involvement of these regions and to clarify their specific role in the pitch-labeling process.

## Author Contributions

S.L, M.G., C.K., and L.J. designed research; M.G. performed research; M.G., and S.S. analyzed data; M.G., S.L., S.S., C.K., and L.J. wrote the paper.

## Conflicts of Interest

The authors declare no conflict of interest.

## Funding sources

This work was funded by the Swiss National Science Foundation (SNSF), grant number 320030_163149 to LJ.

## Acknowledgements

We are particularly grateful to Stefan Elmer for his support. His research has inspired this work greatly. We also wish to thank our research interns and research assistants Anna Speckert, Chantal Oderbolz, Fabian Demuth, Florence Bernays, Isabel Hotz, Joëlle Albrecht, Kathrin Baur, Laura Keller, Melek Haçan, Nicole Hedinger, Pascal Misala, Petra Meier, Piyush Rauch, Sarah Appenzeller, Tenzin Dotschung, Valerie Hungerbühler, Vanessa Vallesi, and Vivienne Kunz for their invaluable assistance with data collection. We thank Christian Brauchli and Anja Burkhard for their contributions within the larger project on absolute pitch.

## Notes

### Competing Interest Statement

The authors have declared no competing interest.

